# The Role of Negatively Charged Groups in Antimicrobial Cationic Peptide Mimics: Insights into Membrane Interactions

**DOI:** 10.1101/2024.12.02.626308

**Authors:** Roni Saiba, Ananya Debnath, Satyavani Vemparala

**Affiliations:** The Institute of Mathematical Sciences, C.I.T. Campus,Taramani, Chennai 600113, India; Homi Bhabha National Institute, Training School Complex, Anushakti Nagar, Mumbai, 400094, India; Department of Chemistry, Indian Institute of Technology Jodhpur, Karwad, Rajasthan, 342037 India

## Abstract

In this study, we explore cationic amphiphilic methacrylate copolymers incorporating both positively charged AEMA and negatively charged PAMA functional groups, focusing on their interactions with bacterial membranes. Aggregation studies reveal that electrostatic interactions drive the formation of stable polymer aggregates, with block copolymers forming micelle-like structures and random copolymers exhibiting a more uniform distribution. These ternary polymers preferentially interact with deep lipid packing defects in bacterial membranes, stabilizing and expanding these defects, while shallow defects remain largely unaffected due to the unfavorable interaction of anionic groups with lipid headgroups. The role of interfacial water is also critical, as hydration layers surrounding anionic groups shield them from electrostatic repulsion, enabling deeper penetration into the membrane. Comparative analyses highlight the advantages of anionic-containing polymers over previously studied polar-containing systems, which predominantly engage shallow defects and exhibit limited structural adaptability near membranes. These findings underscore the role of anionic residues in enabling adaptable AMP conformations, enhanced membrane engagement, and effective disruption mechanisms, providing valuable insights for the design of biomimetic antimicrobial polymers incorporating different functional groups.

## I. INTRODUCTION

Antimicrobial resistance (AMR) poses a critical challenge in healthcare, driven by the increasing ability of bacteria to resist antibiotics. Contributing factors include natural selection and the overuse of antibiotics, resulting in the emergence of multidrug-resistant bacterial strains. The global rise of AMR is approaching a medical crisis, with profound implications for public health [1]. In response, antimicrobial peptides (AMPeptides), naturally occurring in all life forms as primary defenses against bacterial infections, are being explored as potential alternatives to traditional antibiotics. These peptides typically disrupt bacterial membranes through mechanisms such as pore formation and membrane rupture. However, large-scale isolation and purification of AMPeptides remain substantial challenges. To address these limitations, synthetic antimicrobial polymers (AMPs) have been developed using polymer chemistry. These AMPs, based on backbones such as methacrylate, nylon, and norbornene, mimic the functionality of natural AMPeptides and demonstrate particular efficacy against negatively charged bacterial membranes while sparing zwitterionic eukaryotic membranes [2–5].

Antimicrobial peptides are categorized by their monomer charge composition, chemical groups, and structural properties, such as rigidity. Rigid AMPs, like magainin-II, exhibit hydrophilic surfaces that effectively integrate into bacterial membranes. Conversely, flexible AMPs, despite structural differences, can also form amphiphilic surfaces capable of embedding into bacterial membranes [6, 7]. The interaction between AMPs and membranes begins with long-range electrostatic attraction, followed by short-range interactions that facilitate membrane integration. The dynamic lipid-water interface of membranes, along with the presence of non-cylindrical lipids such as phosphatidylethanolamine and polyphosphoinositides, can induce lipid packing defects. These defects expose the hydrophobic core of the membrane, creating sites for interaction with hydrophobic residues of AMPs [8–11]. Traditionally, antimicrobial polymer design has focused on balancing cationic charge and hydrophobicity. Recent advances, however, have expanded this approach to include diverse polymer structures and compositions, such as block copolymers, branched polymers, and polymer assemblies. While cationic amphiphilic copolymers have shown promise against bacterial infections, they often fail to incorporate the diverse functional groups present in natural AMPs, which are critical for stability and specificity toward bacterial membranes. Binary polymers, consisting of cationic and hydrophobic groups, are effective but limited in their ability to modulate the physical and chemical properties of AMPs. To overcome this limitation, research has focused on incorporating functional groups such as tryptophan, guanidine, tyrosine, and hydroxyl groups into the design of antimicrobial copolymers. These efforts have resulted in ternary polymers, which incorporate polar residues alongside binary compnents. Ternary polymers are designed to mimic the amphiphilic *α*-helical structures found in natural AMPs, featuring hydrophobic and hydrophilic surfaces composed of charged and polar residues. This design improves the bioavailability of AMPs, as aggregates formed by ternary polymers exhibit weaker interactions compared to binary polymer aggregates [7, 12].

In this study, we aim to further mimic the functionality of natural AMPs by incorporating anionic groups as a third component in cationic amphiphilic statistical methacrylate copolymers. Natural AMPs, such as defensin, magainin, lactoferricin, LL-37, and dermcidin, contain both positively and negatively charged amino acids in their sequences, despite their overall positive charge, which ranges from +2 to +11 [13–18]. For example, the human cathelicidin AMP LL-37 comprises 11 cationic residues (6 lysine and 5 arginine) and 5 anionic residues (3 glutamic acid and 2 aspartic acid residues). These negatively charged residues play roles beyond simply balancing the overall charge. They contribute to structural stability, selective targeting of bacterial membranes over host membranes, and interaction with the host immune system. Salt bridges formed between negatively charged residues, such as aspartic acid or glutamic acid, and positively charged residues, such as lysine or arginine, stabilize the 3D structure of AMPs, enhancing their functionality. Research highlights the role of these residues in facilitating the formation of oligomeric structures via internal salt bridges [19–21]. Investigations into the charge and composition of AMPs, such as the 26-residue V13K peptide [22], have underscored the nuanced contributions of negatively charged residues to antimicrobial activity, offering critical insights into the design of effective antibiotic alternatives.

Previous studies [7] have explored ternary statistical methacrylate copolymers composed of cationic ammonium, carboxylic acid, and hydrophobic side chain monomers, focusing on their antimicrobial and hemolytic activities as well as chain conformations. These studies demonstrated that antimicrobial and hemolytic activities are primarily influenced by a net positive charge of +3 or higher, with anionic carboxylate groups playing a secondary role. However, anionic groups significantly modulate polymer conformation, enabling dynamic transitions between compact and extended states through salt bridge formation. This flexibility facilitates functional interactions without permanently fixing the polymer structure, broadening the implications for antimicrobial material design. In the current study, we investigate the role of negatively charged groups in AMPs, specifically their interactions with model bacterial membranes and their exploitation of lipid packing defects. Using a random copolymer with a degree of polymerization of 20, featuring balanced positive, negative, and hydrophobic groups, we examine its behavior in solution and at membrane interfaces. Our findings reveal how these polymers exploit lipid packing defects, stabilizing and remodeling membranes. These results provide valuable insights for designing next-generation AMPs with enhanced antimicrobial efficacy.

## II. MODELS AND METHODS

The AMP is composed of three types of monomers: hydrophobic ethyl methacrylate (EMA), positively charged aminoethyl methacrylate (AEMA), and negatively charged propanoic acid methacrylate (PAMA), yielding a net charge of +4 per polymer. The chemical structures of the monomers and the polymer properties are illustrated in Figure 1. All-atom molecular dynamics (MD) simulations were performed using the TIP3P water model [23]. Sicompares the survival probability mulations were conducted with NAMD 2.10 [24] and employed the CHARMM36 force field [25]. The systems were equilibrated and run in the isothermal-isobaric (NPT) ensemble, with hydrogen bonds constrained using the SHAKE algorithm. A timestep of 2 fs was used, and trajectory data was recorded at 40 ps intervals for analysis.

**FIG 1.**
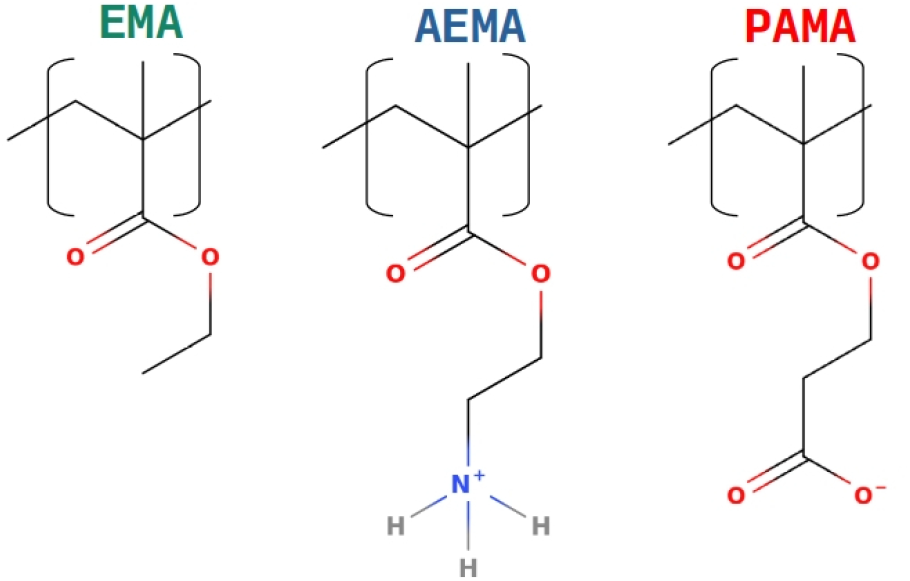
The ternary polymer is composed of hydrophobic ethyl methacrylate (EMA), positively charged aminoethyl methacrylate (AEMA), and negatively charged propanoic acid methacrylate (PAMA).

### A. Polymer aggregate-solution phase

We followed the same procedure, as detailed below, to generate the initial structures and set up simulations for both block and random sequence polymer aggregates (for sequences, see Table I). To create the initial structures, a single AMP polymer was first simulated in aqueous media, neutralized, and equilibrated at 400 K. Conformations were randomly selected from this simulation to serve as starting structures for the aggregate simulations. Twelve such AMP conformations were randomly oriented and positioned in close proximity to one another. The net positive charge of +48 from the AMPs was neutralized using NaCl by adding 116 Na^+^ and 164 Cl^−^ ions, maintaining a salt concentration of 150 mM. The random sequence simulation system contained 127,873 atoms, while the block sequence system contained 126,935 atoms. The temperature was maintained at 310 K with a collision frequency (*γ*) of 5 ps^−1^, and the pressure was held constant at 1 atm using the Langevin piston method. The simulation box size for both systems was approximately 100Å × 110Å × 110Å. Electrostatic interactions were calculated using the Particle Mesh Ewald (PME) method [26], with a cutoff for non-bonded interactions set at 12 Å and smoothing starting at 10 Å. Hydrogen atoms were constrained using the SHAKE algorithm, and a timestep of 2 fs was applied. The systems were first energy-minimized using the conjugate gradient method, followed by 5 ns equilibration in the NVT ensemble. Subsequently, the systems were subjected to NPT simulations. The random sequence polymer system was simulated for approximately 470 ns, while the block sequence polymer system was simulated for approximately 370 ns.

**TABLE I.** Details of AMPs used for simulation. D_*p*_ - degree of polymerization.

| $D_p$ | AEMA | EMA | PAMA | Polymer sequence |
| --- | --- | --- | --- | --- |
| 20 | 8 | 8 | 4 | AEAAEPEAEPEAAEPEAPEA |
| 20 | 8 | 8 | 4 | AAAAAAAAEEEEEEEEPPPP |

**TABLE II.**
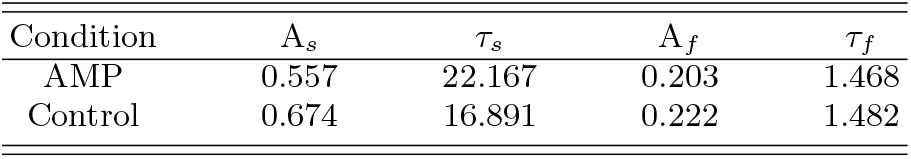
Details of coefficients from biexponential fits.

### B. Multiple polymer-membrane systems

A model Gram-negative bacterial membrane was constructed to investigate the interaction of random sequence AMPs with the membrane. Each leaflet of the membrane comprised 128 lipids, with 98 being POPE residues and 30 POPG residues. Four AMPs were positioned near the membrane surface, resulting in a system with a net charge of -60. This charge was neutralized using NaCl by adding 94 Na^+^ and 34 Cl^−^ ions while maintaining a salt concentration of 150 mM. To enhance statistical reliability, two independent simulations with different initial conditions, referred to as AMP-1 and AMP-2, were conducted for the same system (See SI Figure S1 for snapshots of initial systems). In both setups, four randomly selected equilibrium conformations of AMPs were placed along the membrane normal in the aqueous phase. Simulation parameters for the AMPs were based on previously established methods [12]. The simulation box size for both systems was approximately 90Å × 90Å × 90Å, with the systems containing 67,442 and 83,074 atoms, respectively. The temperature was maintained at 310 K with a collision frequency of 5 ps^−1^, and the pressure was kept constant at 1 atm using the Langevin piston method. Electrostatic interactions were calculated using the Particle Mesh Ewald (PME) method, with a cut-off for non-bonded interactions set at 12 Å and smoothing starting at 10 Å. Both systems were energy-minimized using the conjugate gradient method for 1,000 steps, followed by 5 ns of equilibration in the NVT ensemble. After equilibration, the systems were simulated in the NPT ensemble. The AMP-1 and AMP-2 simulations were run for 730 ns and 1,064 ns, respectively, providing replica simulations to achieve better statistical averages.

### C. Interfacial water analysis

To analyze the properties of interfacial water, 5 ns long NVT simulations were conducted for both AMP-membrane and membrane-only systems. The simulation box volume was propagated from previously described simulations according to the presence or absence of AMP. Data was collected at a frequency of 0.1 ps. Two key quantities were calculated for interfacial water molecules: residence time distribution and survival probability. To identify interfacial water, we first computed the radial distribution function, defining interfacial waters as those within the first peak of the distribution. Based on SI Figure S6, we set a cutoff of 3 Å from membrane phosphate groups to identify interfacial waters. To analyze the residence time distribution, we tracked the duration over which any water molecule remained within 3 Å of membrane phosphate groups. Then we calculated the stretch of time each water molecule spends within this layer. Finally we represented the data as a histogram. To further quantify hydration dynamics at the water membrane interface, we computed the survival probability of interfacial waters. The survival probability *S*(*t*) of waters in a given layer *d* is expressed as:

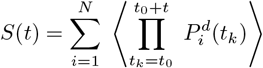

Where 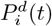 represents the probability that the *i*^*th*^ water molecule is present in layer *d* at time *t*, taking a value of 0 or 1 depending on absence or presence in the layer. This summation across all *N* water molecules is averaged over the initial time origin *t*_0_. The survival probability *S*(*t*) is normalized by the survival probability at time *t* = 0, *S*(0), and is fit to a biexponential function:

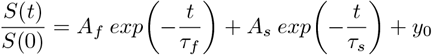

## III. RESULTS

### A. Aggregation dynamics in solution

In this section, we examine how the inclusion of both negatively charged (PAMA) and positively charged (AEMA) functional groups, as well as the sequence of these groups (random vs. block), affects the morphology of polymer aggregates in solution. The ternary polymer system studied here consists of three distinct functional groups, as illustrated in Figure 1. The hydrophobic ethyl methacrylate (EMA) monomers promote aggregation through hydrophobic collapse in aqueous environments, a mechanism commonly observed in amphiphilic polymers and peptides. The positively charged aminoethyl methacrylate (AEMA) monomers facilitate electrostatic interactions with negatively charged counterparts, while the negatively charged propanoic acid methacrylate (PAMA) introduces a crucial counterbalance, modulating the overall electrostatic landscape of the system. This balance allows the system to leverage both hydrophobic and electrostatic forces during aggregation. Unlike binary polymers, which are typically composed of only cationic and hydrophobic monomers and where aggregation is primarily driven by hydrophobic interactions, the addition of negatively charged PAMA reduces the repulsive interactions among cationic groups, enabling the formation of larger, more stable aggregates. This electrostatic balance also prevents the size limitation seen in binary systems, where repulsive forces between similarly charged cationic groups constrain aggregation. Together, this ternary polymer system exhibits a more versatile interplay of interactions, leading to distinct aggregation behaviors based on the sequence and distribution of functional groups.

As detailed in the Methods section, twelve polymers—configured with both random and block sequences (see Table I)—were dispersed in water, and their morphologies were monitored over time. Snapshots of both systems at the end of the simulations are shown in Figure 2. In both cases, complete phase separation occurred, resulting in a single stable aggregate composed of twelve polymers by the end of the simulation. This outcome contrasts with our previous findings [12], where systems containing only hydrophobic and positively charged polymers, or those with a polar functional group, displayed limited aggregation, forming finite bundles rather than undergoing complete phase separation. The current results suggest that favorable electrostatic interactions between the positive and negative functional groups are pivotal in driving the formation of strong aggregates. Such behavior could have significant implications for how individual AMPs or aggregates interact with bacterial membranes, particularly in scenarios where an individual AMP dissociates from the aggregate and partitions into the membrane.

**FIG 2.**
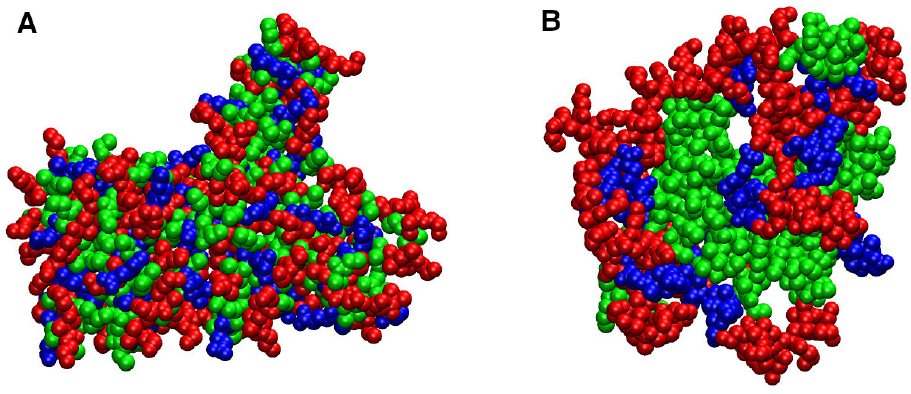
Comparison of the final configurations of random (A) and block (B) sequence aggregates. Red, blue and green denote positive, negative and hydrophobic monomers respectively.

Despite the overall phase separation, notable differences were observed in the morphology and internal arrangement of the polymers between block and random sequences, as illustrated in Figure 2. Block copolymers formed tightly bound micelle-like structures, with hydrophobic groups concentrated at the core and charged groups positioned at the periphery. In contrast, random sequence polymers exhibited a more uniform distribution of functional groups throughout the aggregate. The random sequence polymers form aggregates characterized by an interspersed distribution of hydrophobic, positively charged, and negatively charged monomers. This lack of compartmentalization indicates extensive mixing within the aggregate, likely driven by dynamic inter-molecular salt bridges between AEMA and PAMA groups and the hydrophobic collapse of EMA monomers. In contrast, block sequence polymers exhibit a more organized structure, with EMA monomers forming a compact hydrophobic core, while AEMA and PAMA groups are localized toward the aggregate periphery. This micelle-like structure minimizes unfavorable interactions and reflects intra-polymer organization, which stabilizes the aggregate.

Two quantitative parameters, radius of gyration (*R*_*g*_) and eccentricity (*ϵ*), are monitored as a function of time to characterize the aggregates. The radius of gyration (*R*_*g*_) of an aggregate is given by the root mean square distance of the aggregate atoms measured from its center of mass. The eccentricity *E* is defined as 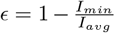, where *I* is the moment of inertia tensor of the aggregate, and *I*_*min*_ and *I*_*avg*_ are the minimum and average of the principal moments of inertia (*I*_1_, *I*_2_, *I*_3_). An eccentricity of *ϵ*= 0 corresponds to a perfectly spherical object, while *ϵ* = 1 represents a rod-like object. The distribution of radius of gyration (*R*_*g*_) and eccentricity (*ϵ*) values for random and block copolymer aggregates, calculated over the last 50 ns of the simulation, are shown in Figure 3. The data in Figure 3(A) shows that the size of the aggregate for block copolymers is larger than that of random copolymers, with a narrower distribution. The data in Figure 3(B) also suggests that the aggregate of block copolymers is significantly more spherical than that of random copolymers. Together, this suggests that the aggregate formed by the random copolymers is more compact yet undergoes dynamic reorganization and has a looser internal structure, likely due to the frustrated arrangement of charged interactions within the aggregate that cannot compartmentalize as effectively as in block copolymer aggregates. This also confirms what is seen visually in Figure 2.

**FIG 3.**
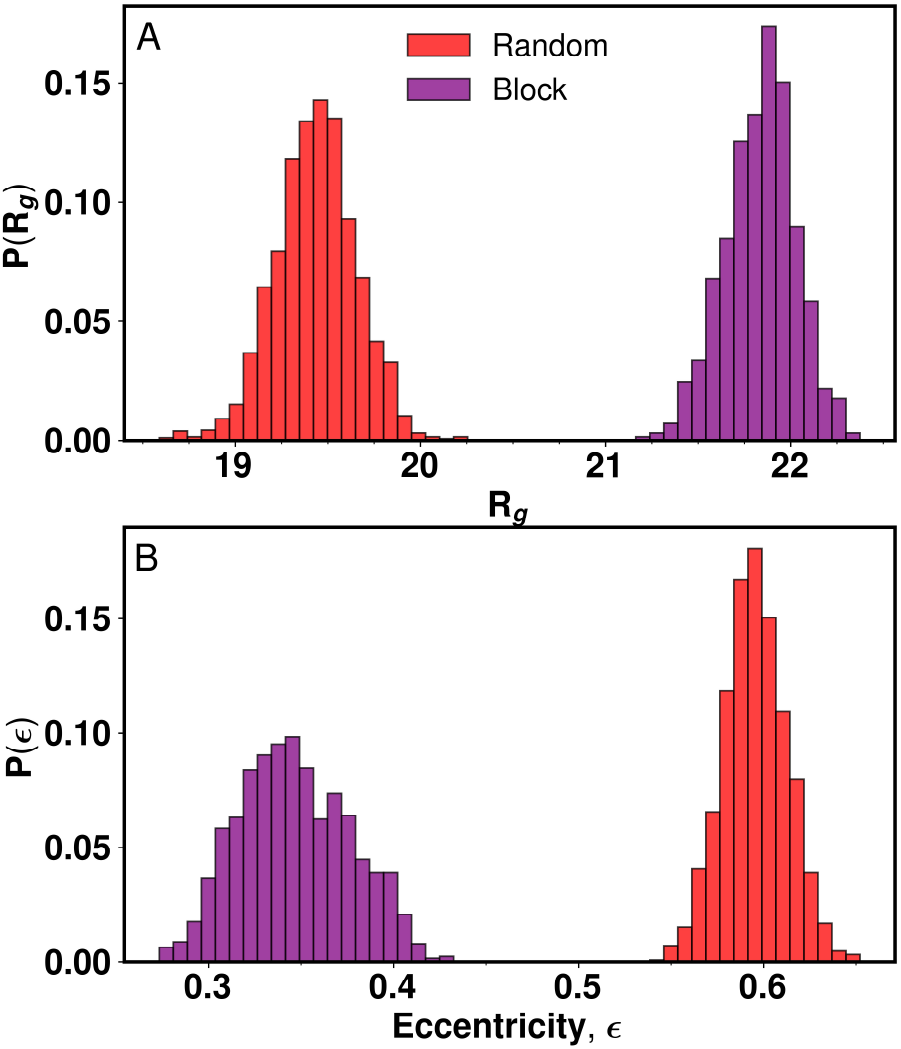
(a) Histogram of distribution of radius of gyration for the block and random aggregates over the last 50 ns of simulation. The random aggregate has lower mean R_*g*_ and variance. Block aggregates show bimodal distribution of R_*g*_ values. (b) Distribution of eccentricity, (*ϵ*) of aggregate with N_*agg*_ = 12 over last 50 ns of simulation.

To understand the nature of electrostatic contacts within the aggregates, we monitored the intra- and interpolymer salt bridge contact fractions over the simulation time for both random and block copolymer aggregates, as shown in Figure 4. Previous work from our group has highlighted the importance of salt bridges in stabilizing single AMP molecules in solution phase [27]. Here, we explore how the salt bridges within the aggregate are influenced by the copolymer sequence, focusing on the interplay between intra-polymer and inter-polymer salt bridges. Salt bridge interactions were defined as a distance of less than 4 Å between the N atom of an amino group and the O atom of a carboxyl group, measured throughout the simulation. In line with the visual representation of aggregate conformations in Figure 2, the temporal evolution of salt bridges suggests that in the long-term limit, inter-polymer contacts for both block and random polymer aggregates are similar. However, as aggregation progresses in the random polymer system, there is a significant reduction in intra-polymer salt bridges and a corresponding increase in inter-polymer salt bridges, indicating enhanced mixing of polymers within the aggregate. In contrast, block copolymers exhibit no significant change in intra-or inter-polymer salt bridges, suggesting a localized compartmentalization of monomers that remains stable throughout the simulation. The presence of strong inter-polymer salt bridges in both block and random copolymer aggregates indicates that electrostatic interactions drive the aggregation of copolymers with positive and negative functional groups. This is in stark contrast to our previous studies [12], where aggregation was driven by strong hydrophobic interactions between polymers containing only positively charged and hydrophobic functional groups. To further investigate the interaction between bulk water and the aggregate, we measured the radial density of water and AMPs from the center of mass of the aggregate (See SI Figure S2). Shells with increasing radii, spaced by 0.1 Å, were used to calculate the density profiles. For the block copolymer, the radial density of water is low at the center, while polymer density is high, indicating the formation of a strong hydrophobic core that expels water. In contrast, the random copolymer aggregate shows higher water density at the center, with maximum polymer density at approximately 20 Å from the center. These results indicate that both the presence and sequence of functional groups along the polymer backbone significantly influence polymer conformations within the aggregate and the aggregate morphology. Since the antimicrobial mechanism of these biomimetic polymers likely depends on their conformation and their release from aggregate into the membrane, these insights into composition and sequence are valuable for advancing AMP design. Our next objectives are to (a) probe the polymer-membrane interactions, (b) analyze polymer conformation and its evolution within a membrane environment compared to solution, and (c) examine any effects the polymer may have on membrane structural properties, particularly on lipid packing defects.

**FIG 4.**
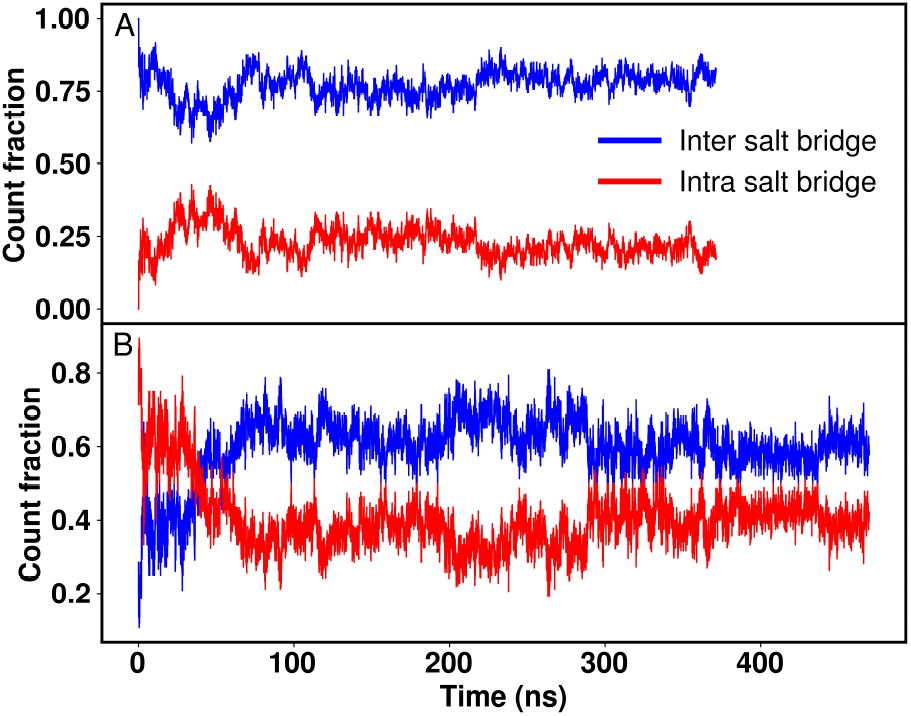
Fraction of intra and interpolymeric salt bridges over the duration of simulation. A - Block sequence aggregate. B - Random sequence aggregate. The fraction of inter polymeric salt bridges increases over time for random sequence but remains more or less constant in case of block sequence aggregates.

### B. Polymer partitioning in model bacterial membranes

In this section, we examine how polymer conformation evolves during partitioning into the membrane and how AMP interactions impact bilayer structure. Facial amphiphilicity is a defining feature of many natural antimicrobial peptides (AMPs), such as LL-37 and magainin, which enables them to selectively target bacterial membranes while sparing host cells [28–32]. This structural characteristic is achieved by organizing hydrophilic and hydrophobic residues on opposite faces of the peptide, creating an amphiphilic surface essential for membrane interactions. Previous simulation studies have shown that binary AMPs, containing only hydrophobic and positively charged (cationic) functional groups, can adopt facial amphiphilicity upon interacting with cell membranes, even when the polymers themselves lack a defined secondary structure [33–37]. However, incorporating polar functional groups alongside hydrophobic and cationic groups has been observed to disrupt facial amphiphilicity, causing AMPs to adopt a more globular conformation instead [7]. Given that the present AMP includes both positively charged AEMA and negatively charged PAMA residues, it is of particular interest to observe the conformations that these membrane-interacting AMPs assume, especially as the model bacterial membrane contains negatively charged lipids.

Snapshots from the simulations reveal that the anionic-containing polymer aggregates exhibit a pronounced tendency for reorganization and interfacial adaptation, forming linear, facially amphiphilic structures upon interacting with bacterial membranes. The interaction of four AMPs with the model bacterial membrane under initial condition 1 is illustrated in SI Figure S3, showing varying degrees of membrane partitioning among the AMPs. By the end of the 1-microsecond simulation, the individual polymers exhibit a range of conformations. In SI Figure S4, the distribution of *R*_*g*_ values for all AMPs in solution-only and membrane simulations are compared, with data computed over both membrane simulation replicas and averaged over the last 100 ns. The results clearly indicate that near the membrane, AMPs tend to adopt more linear and extended conformations not accessible in solution alone. Furthermore, when not adopting extended conformations, AMPs near the membrane show a greater tendency toward compact structures than in the solution phase. This highlights the influence of environment and neighboring AMPs on the conformational landscape. We selected a single AMP that adopts a fully facially amphiphilic conformation during membrane interaction, with snapshots at various simulation timestamps shown in Figure 5. Despite the repulsion between its negatively charged residues and the negatively charged POPG lipids in the bacterial membrane, the AMP notably adopts an extended facially amphiphilic structure, though over longer timescales than its binary counterpart [7, 35]. The snapshots show that one cationic group anchors to the membrane, initiating partitioning. However, the negatively charged groups favor the solution phase, creating a balance between this tendency and favorable interactions between cationic AEMA and POPG, as well as hydrophobic interactions, which likely extends the timescale for complete partitioning.

**FIG 5.**
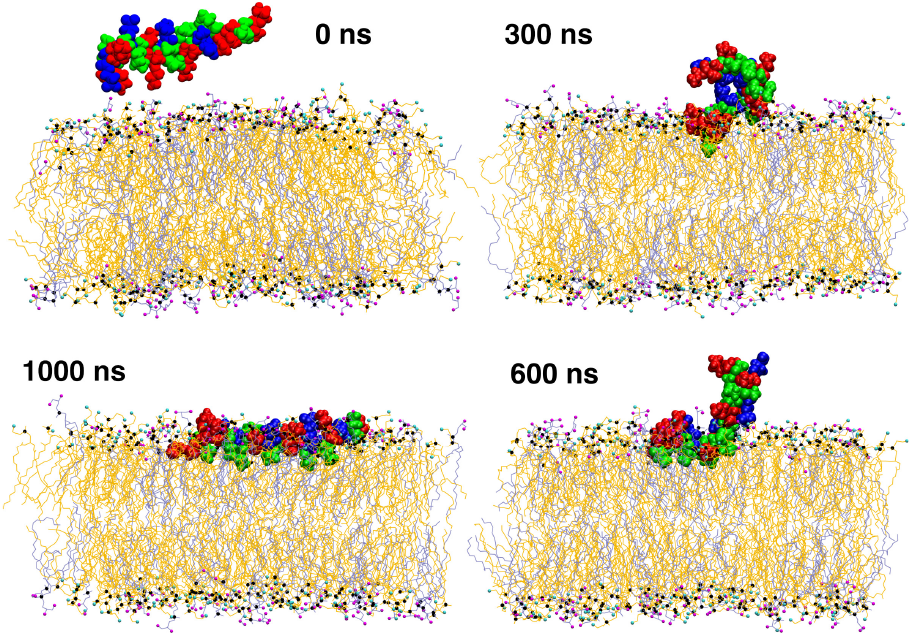
Snapshots of representative AMP and membrane at different time points. Cationic, anionic and hydrophobic groups of the AMP are denoted red, blue and green Van der Waal shpheres respectively. POPE residues are shown as orange lines whereas POPG residues are coloured blue. Phosphate atoms of lipids is represented by black Van der Waal spheres. Terminal nitrogen and oxygen groups of lipids are coloured as cyan and magenta Van der Waal spheres respectively.

The spatial distribution of AMP functional groups along the membrane normal is calculated through *z*-density profiles (averaged over the last 100 ns) and shown in SI Figure S5. The cationic AEMA groups align closely with the phosphate headgroups, consistent with previous simulations of other AMP models containing AEMA. The negatively charged PAMA groups show a strong preference for the solution phase due to repulsive electrostatic interactions with POPG lipids. Hydrophobic EMA groups exhibit a bimodal distribution, indicating an affinity for lipid tails, but the sequence, particularly the presence of PAMA groups, positions them above the headgroups. This suggests that partitioned polymers likely adopt a conformation straddling the solution phase, headgroups, and upper lipid tail regions. To examine how AMP partitioning affects the lateral organization of POPE and POPG lipids, we computed the radial distribution function (*g*(*r*)) of phosphates for POPE and POPG lipids in both membrane leaflets, comparing these with control membranes lacking AMPs (Figure 6). The data indicate a significantly higher accumulation of POPG-POPG lipid molecules in the upper membrane leaflet, where the AMPs partition (evidenced by the increased first peak height in *g*(*r*)), compared to the lower leaflet and control system. Similar POPG clustering in the presence of AMPs was observed in previous studies with binary AMPs [35]. The effects of AMP partitioning on the membrane reorganization, if any, is also probed via computing the lateral membrane thickness distribution in the absence and presence. The data is shown in Figure 7 and is averaged over last 100 ns of membrane simulation 1. The 2-D thickness map is calculated using MEMBPLUGIN [38] extension in VMD by interpolating the distance between the phosphate groups of the upper and lower leaflet into the orthogonal grid in the X-Y plane of the bilayer with a grid spacing of 2Å.

**FIG 6.**
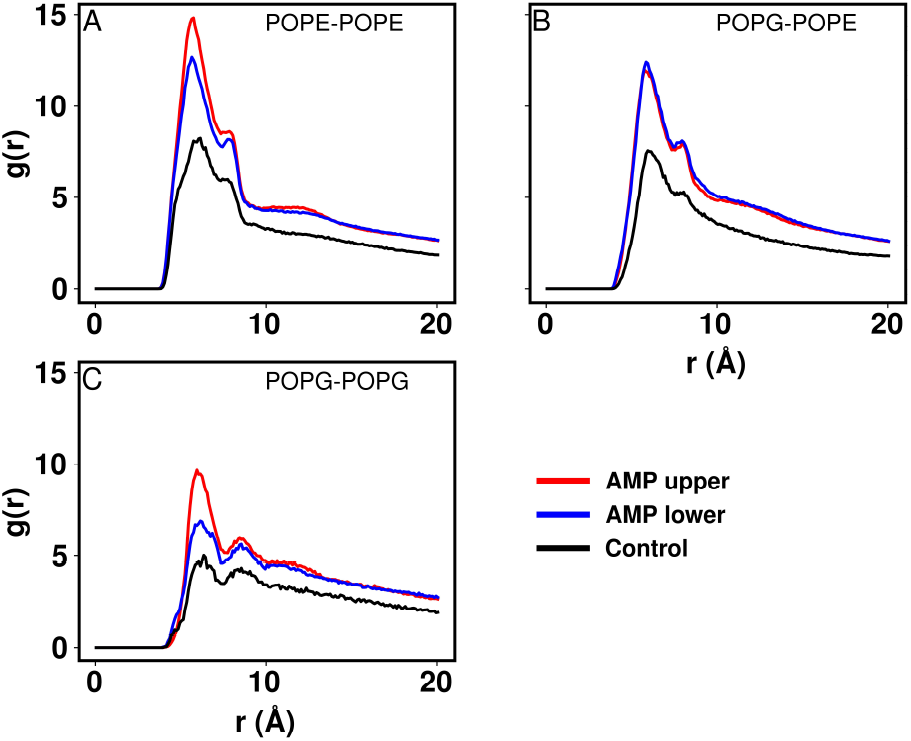
Radial distribution plots of lipid phosphates measured over the last 500 ns of simulation for AMP-membrane system. Control simulation is for 120 ns. RDF is compared between no AMPoly and AMPoly conditions. A - RDF for POPE wrt POPE, B - RDF of POPG wrt POPG, C - RDF of POPG wrt POPE. Presence of AMP increases POPE - POPE clustering.

**FIG 7.**
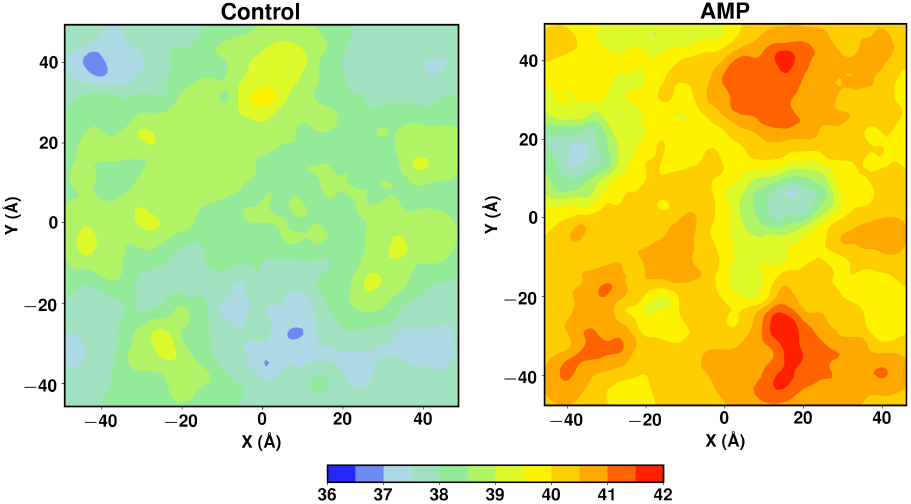
Comparison of membrane thickness, computed over last 100 ns. with AMP and without AMP (control).

Average membrane thickness was measured by calculating the difference between average position of P atom in upper and lower leaflets of the membrane. The change in average membrane thickness is higher than previously reported for ternary methacrylate polymers composed of cationic, polar and hydrophobic groups. Further analysis was also done to measure patchwise thickness of the membrane. Compared to the previous study with polar residues, the difference between the maximum and minimum thickness of the membrane is also higher.

### C. Interfacial water

The dynamics and structure of water molecules at lipid interfaces have been extensively studied [39–42]. In this work, we specifically examine how the interfacial hydration layer near the bilayer, in the presence of AMPs, facilitates membrane partitioning and whether AMPs alter hydration dynamics. The residence time distribution Figure 8(A) indicates a significantly longer residence time for interfacial waters in the presence of AMPs when compared to control. We observed the long lived water molecules individually and these were sandwiched between the AMP and the upper leaflet of membrane bilayer. Figure 8(B) compares the survival probability of interfacial water in AMP and control cases, calculated over a continuous 40 ps window with a 1 fs sampling frequency from the 5 ns trajectory. Fitting with the biexponential model reveals that the fast decay in survival probability remains largely unchanged; however, the presence of AMPs significantly increases the overall survival probability of water molecules. These findings strongly suggest that the hydration layer near the membrane head-groups exhibits altered behavior in the presence of AMPs.

**FIG 8.**
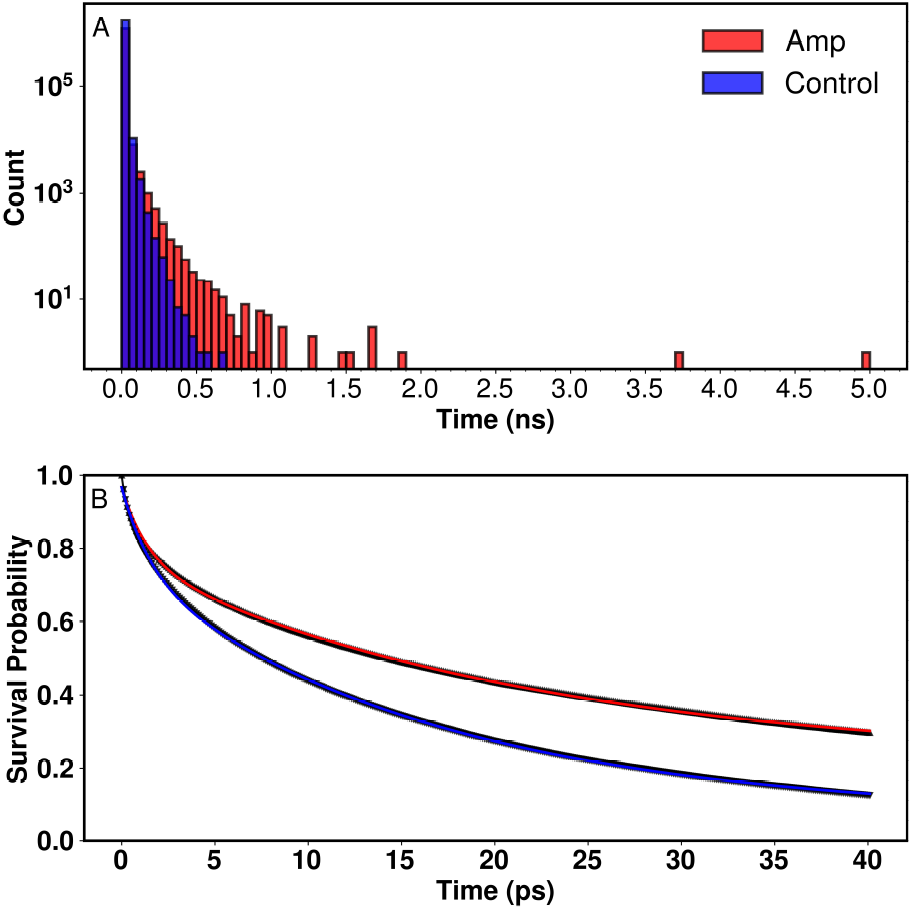
Comparison of A-residence time and B-survival probalility of interfacial water with AMP and lipid control. Interfacial waters are selected as water within 3 Å of mean upper leaflet phosphate position. Addition of AMP increases both residence time and survival probability of interfacial water.

To further explore interfacial water’s role in AMP-membrane interactions, we examined how water molecules are distributed around different AMP functional groups, aiming to understand how interfacial water may shield the negatively charged PAMA groups from the anionic POPG lipids. The *g*(*r*) data for AMP functional groups and water, presented in SI Figure S6(A), shows an increased concentration of water surrounding the negatively charged PAMA groups, consistent with earlier *z*-density profiles of AMPs along the membrane normal. This distribution may help explain how ternary AMPs can adopt facially amphiphilic conformations despite containing negatively charged PAMA groups, as these groups are effectively shielded from direct contact with lipid headgroups by long-residing water molecules. Additionally, the *g*(*r*) data in SI Figure S6(B,C) shows a strong tendency for positively charged AEMA groups to interact with POPG and POPE lipid molecules, while interactions between PAMA and POPG and POPE are minimal. Thus, interfacial water likely plays a crucial role in mitigating unfavorable electrostatic repulsion between PAMA and POPG molecules, facilitating AMP adoption of linear, facially amphiphilic conformations.

### D. Interaction between AMP and lipid packing defects

Thermal fluctuations at the lipid-water interface of a membrane can lead to transient lipid packing defects, which are randomly formed. Many viral peptides and AMPeptides exploit these defects to interact with membranes. Packing defects are classified as shallow or deep, depending on their depth relative to the glycerol C atom of the lipid. Our analysis shows that the AMP interacts specifically with deep defects, as evidenced by an increase in the total area and size of deep defect sites upon AMP addition, compared to control simulations. Interestingly, no significant change was observed in shallow defects, indicating a preferential binding of the AMP to deep defects. The lipid-water interface is highly dynamic, with fluctuations that expose hydrophobic regions of lipid hydrocarbon tails on nanometer length scales and picosecond timescales. These exposed hydrophobic sites are critical for AMP binding, as shown in previous AM-Peptide studies where deep defects facilitated membrane partitioning. Prior studies with ternary AMPs containing positive, negative, and polar residues also demonstrated a skew in the deep defect size distribution towards larger values, while shallow defect distribution remained unchanged [9]. We employed the PACKMEM software package [43] to analyze AMP-lipid defect interactions. Defects deeper than the C2 atom of the lipid were classified as deep, while those above were considered shallow. Defects were characterized at each frame over the initial and final 300 ns of the simulation, with frames separated by 120 ps, exceeding the average defect lifetime of 20 ps. Analyses were conducted for both the AMP-proximal (upper) and AMP-distal (lower) membrane leaflets. Individual lipid defect areas were calculated for both leaflets (see Figure 9). Results indicate a significant increase in larger deep defect sites in the upper leaflet in the presence of AMP, while the lower leaflet’s deep defect size distribution remained consistent between the initial and final simulation stages. The shallow defect distribution showed no notable change with AMP addition. Further comparison of total defect areas revealed a marked increase in deep defect area in the upper leaflet, with minimal change in the lower leaflet as seen in Figure 10. Concomitantly, the shallow defect area slightly decreased in the upper leaflet, with no change in the lower. Collectively, these findings suggest that ternary AMPs interact with membranes primarily via deep defect sites.

**FIG 9.**
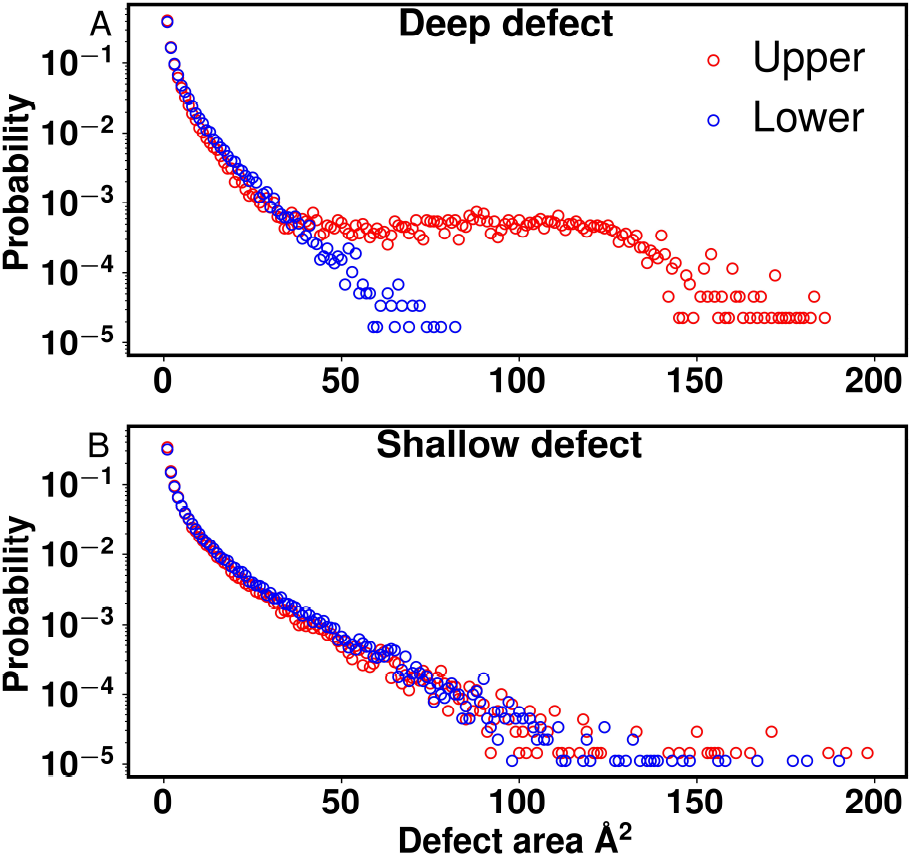
Comparison of deep and shallow defect area distributions in upper (red) and lower (blue) leaflets of the membrane for final 300 ns of simulation. Addition of AMP markedly increases the prevalence of larger deep defects whereas there is no pronounced effect for shallow defects.

**FIG 10.**
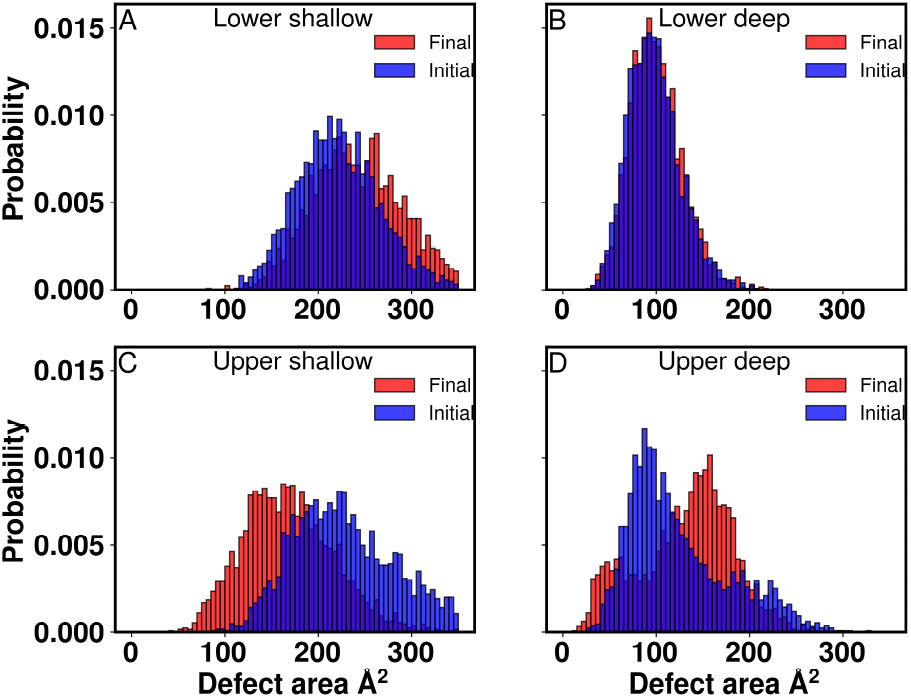
Histograms of total area of different defect types over the course of initial and final 300 ns of simulation. Deep defect sites increase in the upper leaflet (A) while the distribution is more or less unchanged in lower leaflet. Shallow defect area distribution in upper leaflet shows a slight decrease (B) while it is similar in lower leaflet.

Since the total deep defect area in the upper leaflet increased with AMP addition, we investigated whether this was due to the formation of new deep defects or the fusion and stabilization of existing ones. We calculated the number of deep and shallow defects in each leaflet over the initial and final 300 ns of the simulation (See SI figure S7). The addition of AMP led to a decrease in the count of deep defects, along with a concurrent reduction in shallow defects in the upper leaflet, while no such change was observed in the lower leaflet. Given the observed increase in total deep defect area in the presence of AMP, we propose that AMPs stabilize deep defects on the membrane surface, facilitating their fusion and growth. This stabilization likely contributes to the observed decrease in the number and area of shallow defects in the upper leaflet. These findings further support the conclusion that ternary negative AMPs interact with the membrane primarily via deep defects. This selective amplification of deep defects can be attributed to the chemical composition of the polymer. The positively charged AEMA groups strongly associate with the negatively charged POPG lipids, as shown by the pronounced peak in the radial distribution function (RDF) at ∼ 4–5Å (See SI Figure S6). This interaction anchors the polymer to the membrane surface, allowing the hydrophobic EMA groups to penetrate deeper into the bilayer and engage with hydrophobic regions exposed in deep defects. This anchoring facilitates a mechanism where the polymer stabilizes and expands deep defects while leaving shallow defects largely unaffected. The negligible interaction between the negatively charged PAMA groups and the POPG headgroups, as indicated by the flat RDF profile, further explains the lack of shallow defect engagement. Electrostatic repulsion between the PAMA and POPG groups drives the polymer away from headgroup-dominated regions, bypassing shallow defects and targeting deeper regions within the membrane. This behavior contrasts sharply with the findings from our earlier study [44] on ternary polymers with polar groups instead of anionic groups. In those systems, the polar groups could form weakly favorable interactions with lipid head-groups, leading to a predominant engagement with shallow defects. Here, the inclusion of PAMA not only shifts the interaction profile but also enables a unique mode of action where interfacial water plays a crucial role.

## IV. DISCUSSION

This study investigates biomimetic ternary methacrylate antimicrobial polymers (AMPs), focusing on the role of incorporating positive and negative charges alongside hydrophobic groups. Antimicrobial peptides (AMPeptides), naturally found across all domains of life, are key components of innate immunity, combating bacterial infections by disrupting membranes through mechanisms such as pore formation and membrane destabilization. However, challenges in isolating and purifying natural AMPs have prompted the development of synthetic AMP-mimics that replicate their functional properties using polymer chemistry. Synthetic AMPs incorporate diverse functional groups, including hydrophobic, cationic, polar, and anionic moieties, to mimic the structural versatility and targeted activity of natural AMPs. Binary AMPs, composed of cationic and hydrophobic groups, aggregate primarily through hydrophobic collapse, but their antimicrobial activity is often limited by nonspecific interactions with eukaryotic membranes and by size-constrained aggregates due to repulsive forces between like charges. Ternary systems, incorporating polar or anionic groups, aim to overcome these limitations by modulating electrostatic and hydrophobic interactions, achieving greater selectivity and efficiency in bacterial targeting. In natural AMPs such as LL-37, defensins, and dermcidin, negatively charged residues (e.g., glutamic acid and aspartic acid) coexist with cationic residues, contributing to structural stability, specificity, and functionality. These anionic residues form intramolecular salt bridges with cationic residues, stabilizing secondary and tertiary structures critical for dynamic environments like membrane interfaces. They also reduce nonspecific electrostatic interactions with bacterial lipids, preventing excessive aggregation on membrane surfaces. Additionally, anionic residues play immunomodulatory roles by interacting with receptors such as Toll-like receptors (TLRs) and influencing cytokine production, enabling AMPs to act as both antimicrobial agents and regulators of the immune response. Although synthetic AMPs are not intended to mimic these immunological roles, understanding the function of anionic residues in natural AMPs offers valuable insights for designing multifunctional AMP mimics.

The inclusion of negatively charged PAMA groups in this study provides an electrostatic counterbalance to the positively charged AEMA groups, modulating aggregation behavior and facilitating conformational adaptability. This ternary polymer system is distinguished from both binary systems and ternary systems incorporating polar groups, exhibiting unique interactions with bacterial membranes. Our results emphasize the importance of sequence-dependent aggregation behavior in solution. Random sequences, with their mixed organization of functional groups, form more dynamic aggregates with enhanced inter-polymer interactions. These aggregates demonstrate extensive mixing and a lack of compartmentalization, reflecting the dynamic flexibility of natural AMPs. In contrast, block-sequence polymers form more rigid, micelle-like structures where hydrophobic components are sequestered in a core, and charged components are localized at the periphery. These differences in aggregation arise from the interplay of hydropho-bic and electrostatic forces, with random sequences favoring inter-polymer salt bridges and block sequences favoring intra-polymer interactions. The ternary polymer containing cationic (AEMA), hydrophobic (EMA), and anionic (PAMA) groups demonstrated a significant ability to form facially amphiphilic conformations during membrane interactions. This structural adaptability facilitates deeper membrane partitioning and enhanced stabilization of lipid packing defects compared to ternary systems with polar groups studied previously [7]. Polymers containing polar groups exhibited limited aggregation and membrane interaction, primarily associating with shallow lipid packing defects and remaining near the aqueous interface. In contrast, the anionic-containing AMPs in this study preferentially interacted with deep defects, enabling more stable and extended conformations within the membrane. These results align with observations from natural AMPs, where deep defect interactions are critical for selective embedding in bacterial membranes. The dynamic charge balance provided by PAMA and AEMA groups promotes adaptable electrostatic interactions, enhancing the polymers’ ability to adopt facially amphiphilic conformations.

The influence of interfacial water dynamics further highlights the role of anionic groups in modulating membrane interactions. In the anionic-containing polymer system, hydration dynamics mitigate repulsive electrostatic interactions between PAMA and negatively charged POPG lipids. Radial distribution function (RDF) analyses show that water molecules preferentially associate with PAMA groups, forming a hydration layer that shields these groups from direct interaction with the membrane surface. This shielding effect enables the polymers to adopt linear, facially amphiphilic conformations, facilitating deeper partitioning into the lipid bilayer. Consistent with studies on polar-group AMPs, our results demonstrate that ternary AMPs alter interfacial water dynamics, increasing residence time and stability of interfacial water molecules. This altered hydration environment likely supports dynamic membrane engagement and defect stabilization, enhancing antimicrobial efficacy. The aggregation behavior of these polymers in aqueous environments reflects the interplay between hydrophobic and electrostatic interactions. Unlike binary AMPs, which exhibit size-constrained aggregates due to cationic repulsion, the inclusion of PAMA reduces these repulsive forces, enabling larger and more stable aggregates. The radial density profiles reveal that block copolymers form dense hydrophobic cores that exclude water, while random copolymers exhibit higher water density at the core, indicative of looser, dynamically reorganizing aggregates. These differences have important implications for antimicrobial mechanisms, as aggregate morphology and stability influence polymer-membrane interactions, including dissociation and partitioning of individual AMPs.

Finally, our simulation results reveal that membrane partitioning and defect interactions in anionic-group AMPs occur over extended timescales. The anionic groups create a competitive dynamic, counterbalancing the cationic attraction to membrane lipids and leading to gradual, yet deeper, partitioning compared to binary AMPs. The increased total area and size of deep defects in the presence of AMPs indicate that these polymers stabilize and expand existing defects, promoting sustained membrane interaction and structural disruption. Interestingly, while deep defect interactions were amplified, shallow defect sites showed minimal change, suggesting a specific affinity of anionic AMPs for structurally impactful binding sites within the membrane. Collectively, these findings demonstrate the importance of incorporating anionic groups into AMP design. Far from being passive components, anionic groups contribute to structural stability, selective interactions, and enhanced membrane-targeting efficiency. By mimicking features of natural AMPs, such as electrostatic balance and dynamic structural adaptability, anionic-containing polymers provide new strategies for developing effective antimicrobial materials to combat resistant bacterial strains. Future efforts should focus on optimizing the balance of hydrophobic, cationic, and anionic groups and investigating the effects of sequence variations and additional functional modifications to enhance AMP-mimetic performance against antibiotic-resistant bacteria.

## V. ACKNOWLEDGMENTS

All the simulations in this work have been carried out on clusters Nandadevi and Kamet at The Institute of Mathematical Sciences, Chennai, India.

## References

[1] C. J. Murray, K. S. Ikuta, F. Sharara, L. Swetschinski, G. R. Aguilar, A. Gray, C. Han, C. Bisignano, P. Rao, E. Wool, et al., The Lancet 399, 629 (2022).

[2] Z. M. Al Badri, A. Som, S. Lyon, C. F. Nelson, K. Nusslein, and G. N. Tew, Biomacromolecules 9, 2805 (2008).

[3] K. Kuroda and W. F. DeGrado, Journal of the American Chemical Society 127, 4128 (2005).

[4] M. W. Lee, S. Chakraborty, N. W. Schmidt, R. Murgai, S. H. Gellman, and G. C. Wong, Biochimica et Biophysica Acta (BBA)-Biomembranes 1838, 2269 (2014).

[5] H. Takahashi, I. Sovadinova, K. Yasuhara, S. Vemparala, G. A. Caputo, and K. Kuroda, Wiley Interdisciplinary Reviews: Nanomedicine and Nanobiotechnology 15, e1866 (2023).

[6] F. Siedenbiedel and J. C. Tiller, Polymers 4, 46 (2012).

[7] G. Rani, K. Kuroda, and S. Vemparala, Soft Matter 17, 2090 (2021).

[8] N. van Hilten, K. S. Stroh, and H. J. Risselada, Journal of Chemical Theory and Computation 18, 4503 (2022).

[9] S. Sikdar, M. Banerjee, and S. Vemparala, The Journal of membrane biology 255, 129 (2022).

[10] S. Sikdar, M. Banerjee, and S. Vemparala, Soft Matter 17, 7963 (2021).

[11] M. M. Ouberai, J. Wang, M. J. Swann, C. Galvagnion, T. Guilliams, C. M. Dobson, and M. E. Welland, Journal of Biological Chemistry 288, 20883 (2013).

[12] G. Rani, K. Kuroda, and S. Vemparala, Journal of Physics: Condensed Matter 33, 064003 (2020).

[13] U. H. Dürr, U. Sudheendra, and A. Ramamoorthy, Biochimica et Biophysica Acta (BBA)-Biomembranes 1758, 1408 (2006).

[14] J. Johansson, G. H. Gudmundsson, M. E. Rottenberg, K. D. Berndt, and B. Agerberth, Journal of Biological Chemistry 273, 3718 (1998).

[15] Š. Gruden and N. Poklar Ulrih, International journal of molecular sciences 22, 11264 (2021).

[16] H. J. Min, H. Yun, S. Ji, G. Rajasekaran, J. I. Kim, J.-S. Kim, S. Y. Shin, and C. W. Lee, Scientific Reports 7, 45282 (2017).

[17] M. Yang, C. Zhang, M. Z. Zhang, and S. Zhang, BMC microbiology 18, 1 (2018).

[18] G. Laverty, in Peptides and proteins as biomaterials for tissue regeneration and repair (Elsevier, 2018) pp. 347– 368.

[19] C. Aisenbrey, M. Amaro, P. Pospíšil, M. Hof, and B. Bechinger, Scientific reports 10, 11652 (2020).

[20] T. H. Walther, C. Gottselig, S. L. Grage, M. Wolf, A. V. Vargiu, M. J. Klein, S. Vollmer, S. Prock, M. Hartmann, S. Afonin, et al., Cell 152, 316 (2013).

[21] H. Kim, J. Hsin, Y. Liu, P. R. Selvin, and K. Schulten, Structure 18, 1443 (2010).

[22] Z. Jiang, A. I. Vasil, J. D. Hale, R. E. Hancock, M. L. Vasil, and R. S. Hodges, Peptide Science 90, 369 (2008).

[23] W. L. Jorgensen, J. Chandrasekhar, J. D. Madura, R. W. Impey, and M. L. Klein, The Journal of chemical physics 79, 926 (1983).

[24] J. C. Phillips, D. J. Hardy, J. D. Maia, J. E. Stone, J. V. Ribeiro, R. C. Bernardi, R. Buch, G. Fiorin, J. Hénin,W. Jiang, et al., The Journal of chemical physics 153, 044130 (2020).

[25] J. B. Klauda, R. M. Venable, J. A. Freites, J. W. O’Connor, D. J. Tobias, C. Mondragon-Ramirez, Vorobyov, A. D. MacKerell Jr, and R. W. Pastor, The journal of physical chemistry B 114, 7830 (2010).

[26] U. Essmann, L. Perera, M. L. Berkowitz, T. Darden, H. Lee, and L. G. Pedersen, The Journal of chemical physics 103, 8577 (1995).

[27] R. Bhat, L. L. Foster, G. Rani, S. Vemparala, and K. Kuroda, RSC advances 11, 22044 (2021).

[28] M. Radic and S. Muller, “Ll-37, a multi-faceted amphipathic peptide involved in netosis,” (2022).

[29] K. Zeth and E. Sancho-Vaello, International Journal of Molecular Sciences 22, 5200 (2021).

[30] X. Li, Y. Li, H. Han, D. W. Miller, and G. Wang, Journal of the American Chemical Society 128, 5776 (2006).

[31] E. Sevcsik, G. Pabst, W. Richter, S. Danner, H. Amenitsch, and K. Lohner, Biophysical journal 94, 4688 (2008).

[32] J. Zerweck, E. Strandberg, O. Kukharenko, J. Reichert, J. Bürck, P. Wadhwani, and A. S. Ulrich, Scientific reports 7, 13153 (2017).

[33] E. F. Palermo, S. Vemparala, and K. Kuroda, “Antimicrobial polymers: molecular design as synthetic mimics of host-defense peptides,” in Tailored Polymer Architectures for Pharmaceutical and Biomedical Applications (American Chemical Society, 2013) Chap. 20, pp. 319–330.

[34] E. F. Palermo, S. Vemparala, and K. Kuroda, Biomacromolecules 13, 1632 (2012).

[35] U. Baul, K. Kuroda, and S. Vemparala, The Journal of Chemical Physics 141, 084902 (2014).

[36] M. A. Rahman, M. Bam, E. Luat, M. S. Jui, M. S. Ganewatta, T. Shokfai, M. Nagarkatti, A. W. Decho, and C. Tang, Nature Communications 9, 5231 (2018).

[37] U. Baul and S. Vemparala, Soft Matter 13, 7665 (2017).

[38] R. Guixá-González, I. Rodriguez-Espigares, J.M. Ramírez-Anguita, P. Carrió-Gaspar, H. Martinez-Seara, T. Giorgino, and J. Selent, Bioinformatics 30, 1478 (2014).

[39] A. Debnath, B. Mukherjee, K. Ayappa, P. K. Maiti, and S.-T. Lin, The Journal of Chemical Physics 133 (2010).

[40] S. Malik, S. Karmakar, and A. Debnath, The Journal of Chemical Physics 158 (2023).

[41] J. C. Flanagan, M. L. Valentine, and C. R. Baiz, Accounts of chemical research 53, 1860 (2020).

[42] L. R. Franco, P. Park, H. Chaimovich, K. Coutinho, M. Cuccovia, and F. S. Lima, RSC advances 12, 4573 (2022).

[43] R. Gautier, A. Bacle, M. L. Tiberti, P. F. Fuchs, S. Vanni, and B. Antonny, Biophysical journal 115, 436 (2018).

[44] S. Sikdar, G. Rani, and S. Vemparala, Langmuir 39, 4406 (2023).

