## Supplemental Information for "The Role of Negatively Charged Groups in Antimicrobial Cationic Peptide Mimics: Insights into Membrane Interactions"

(Dated: 2 December 2024)

[a] Electronic mail:

[b] Electronic mail:

[c] Electronic mail:

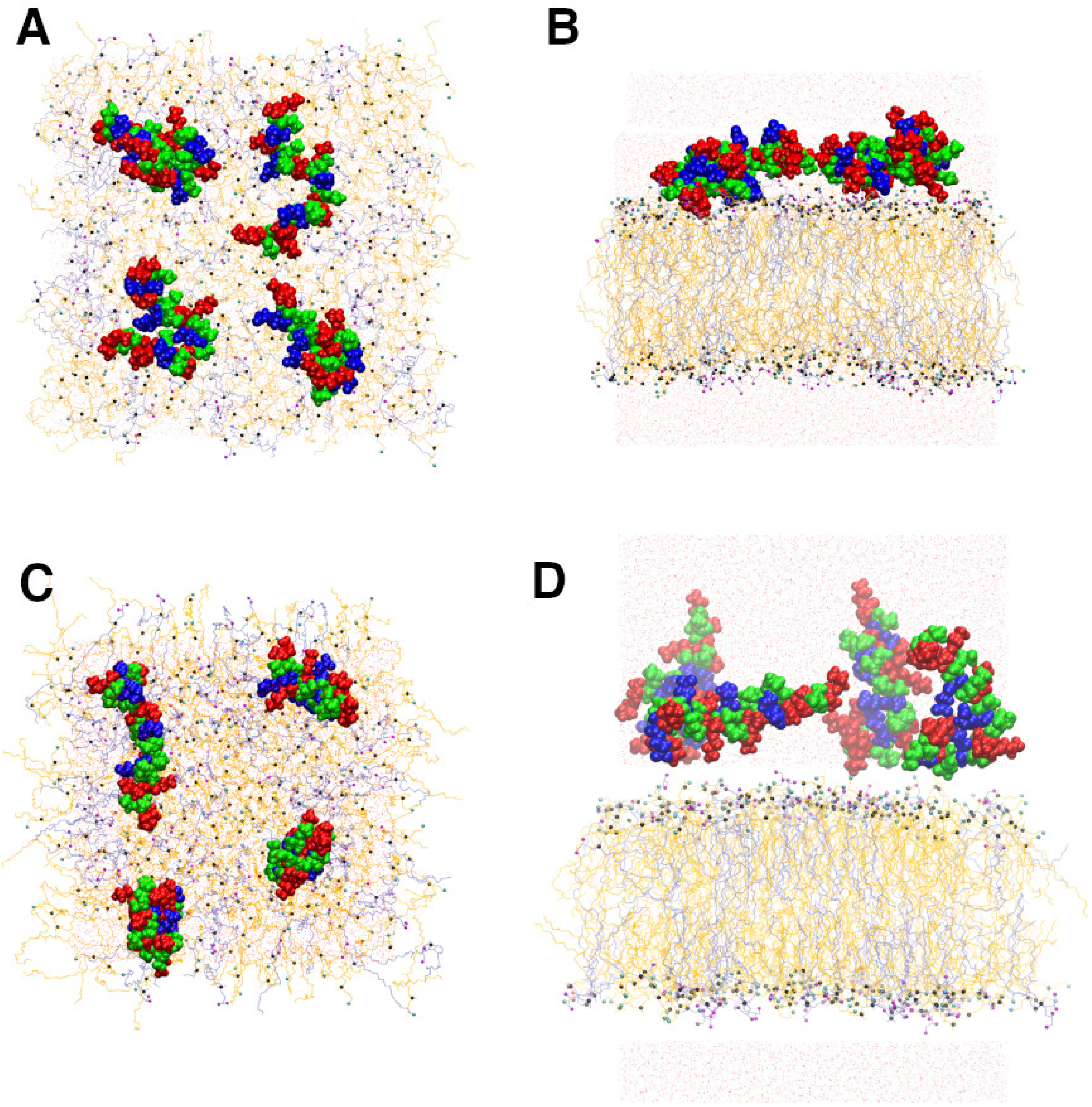


FIG. S1. The initial placement of four AMPs in the solution phase near the model bacterial membrane. A-B for Simulation 1 and C-D for Simulation 2.

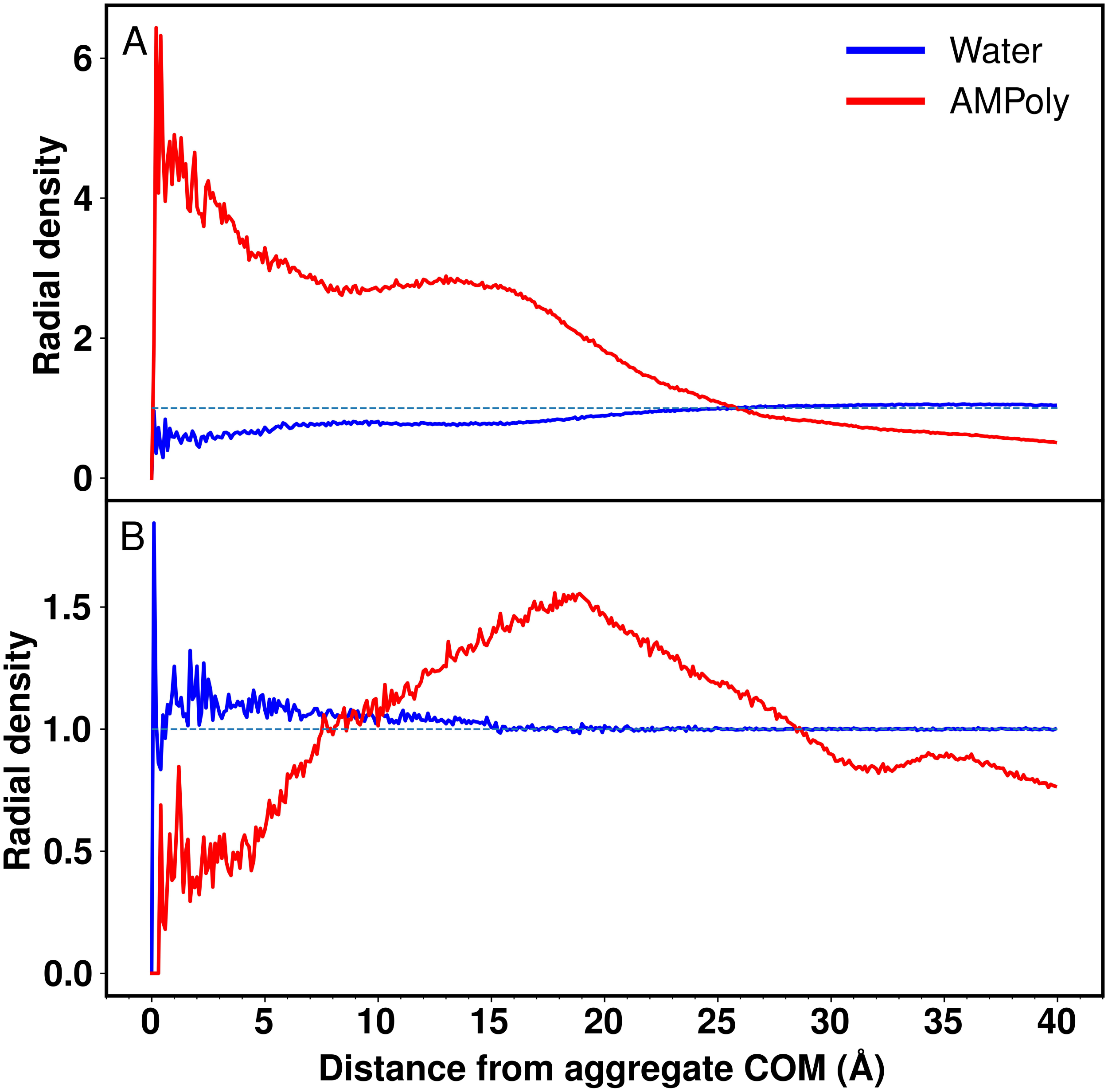


FIG. S2. Radial distribution of water from the center of aggregate (A) Block copolymers (B) Random copolymers

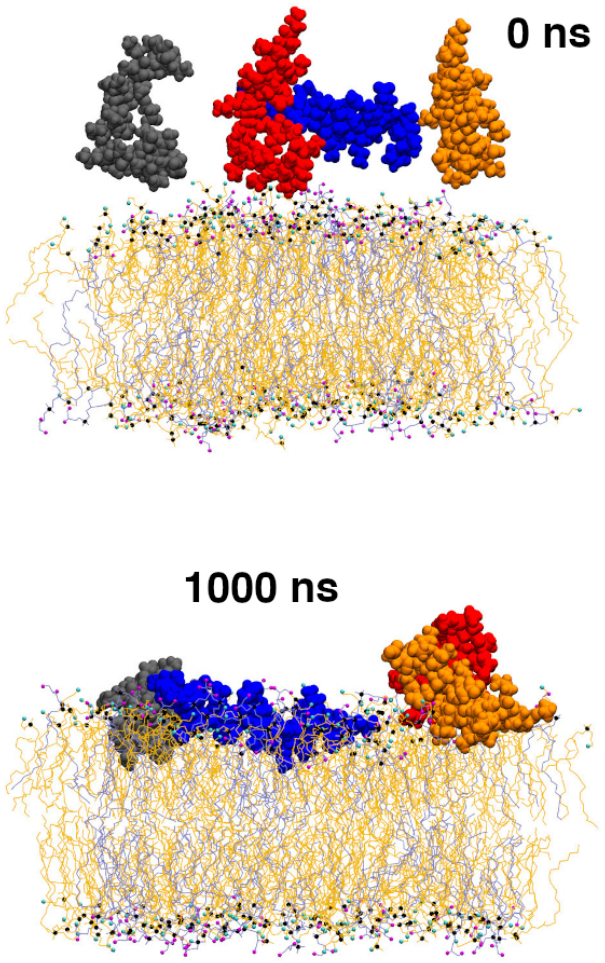


FIG. S3. Snapshots of four AMP and membrane at different time points. The four AMPs are shown in different colours and in Van der Waal spheres respectively. POPE residues are shown as orange lines whereas POPG residues are coloured blue. Phosphate atoms of lipids is represented by black Van der Waal spheres. Terminal nitrogen and oxygen groups of lipids are coloured as cyan and magenta Van der Waal spheres respectively

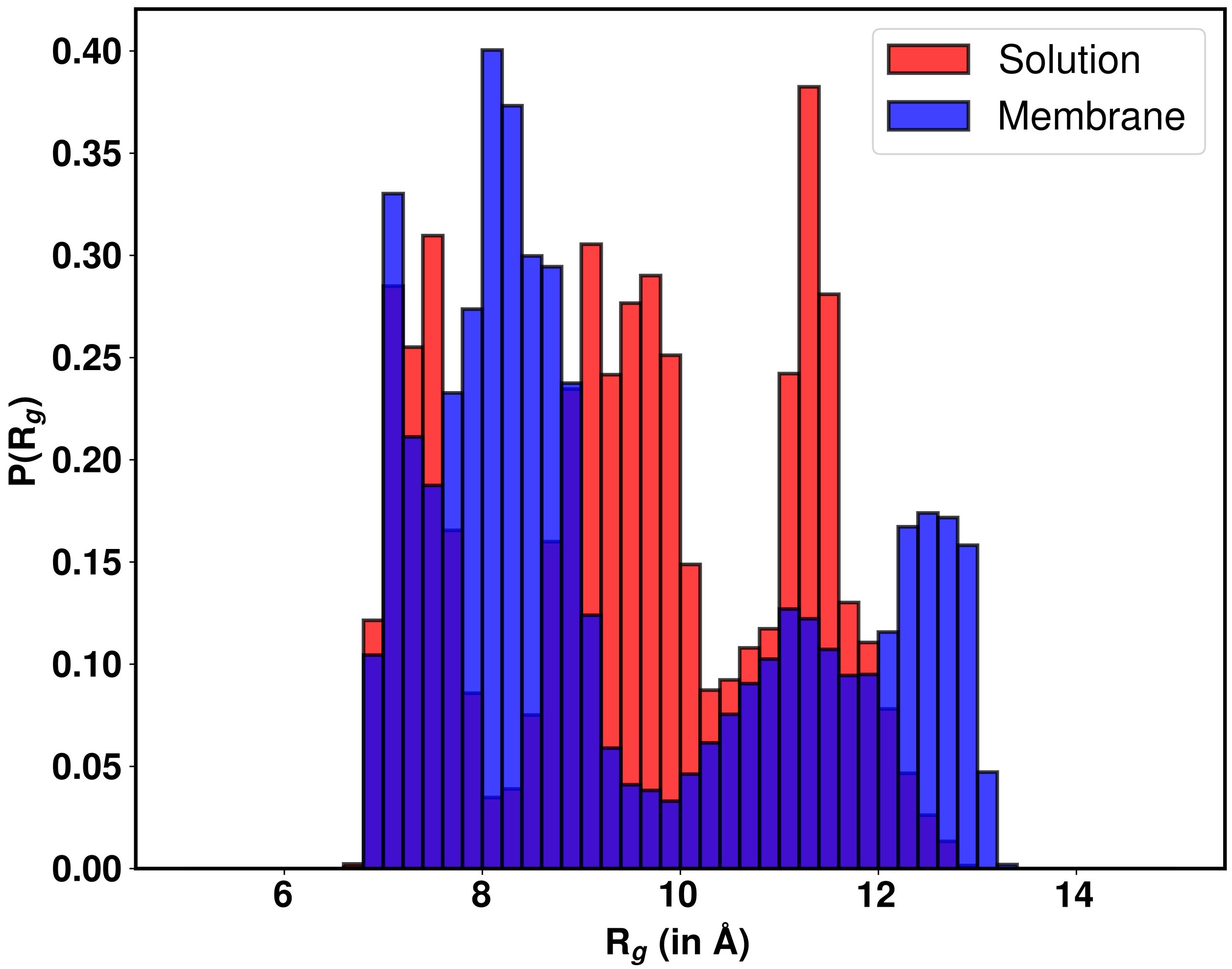


FIG. S4. Distribution of radius of gyration ($R_g$) of individual polymers in solution and in membrane systems. For teh membrane systems, all the polymers in two simulations were considered for the distribution. The data is computed over last 100 ns of all simulations.

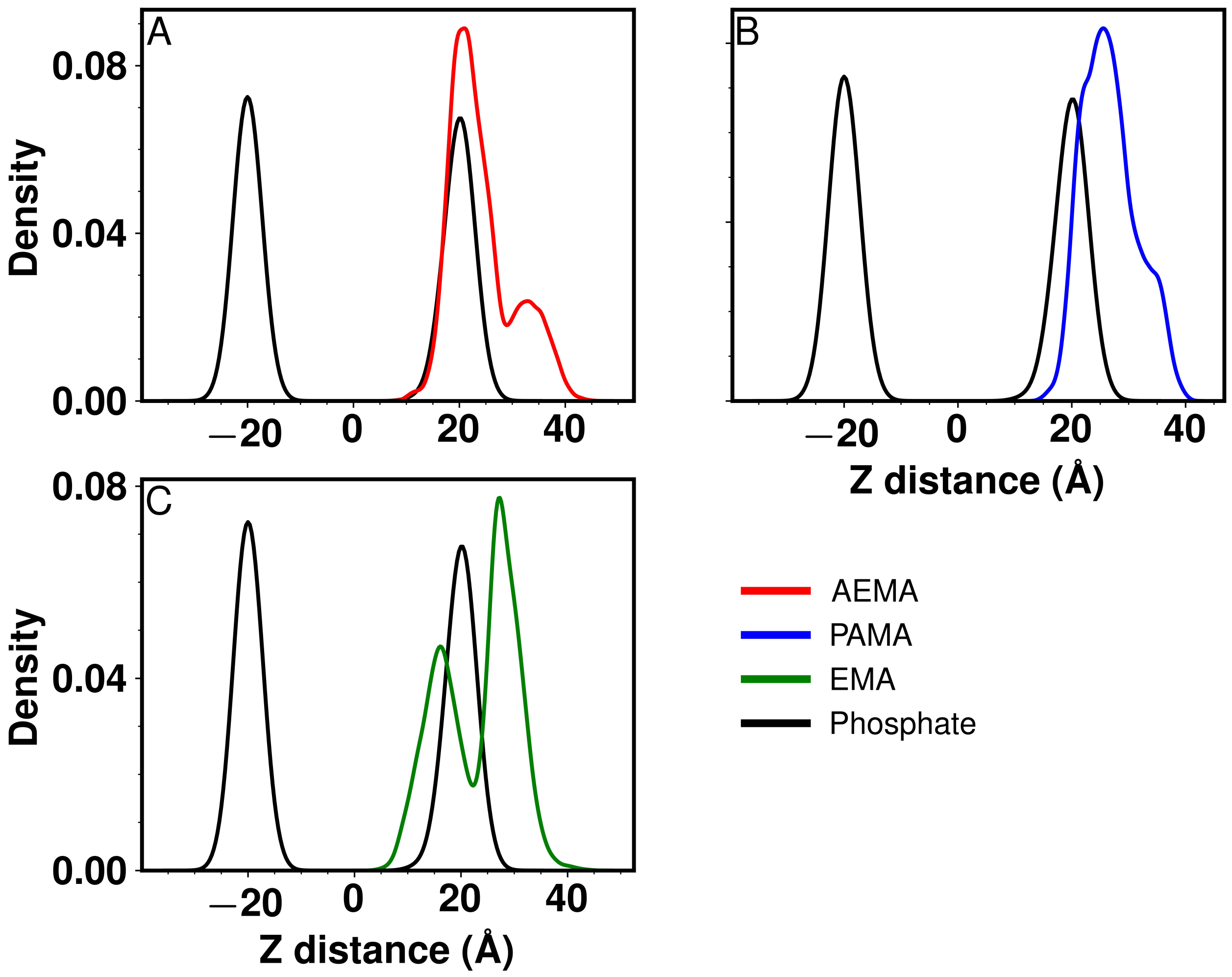


FIG. S5. Density plots of different groups over the last 100 ns of simulation. For the membrane the phosphate and hydrdoxyl groups are shown. The position of amino nitrogen (AEMA), carbonyl oxygen (PAMA) and terminal aliphatic carbon (EMA) is shown for the polymer.

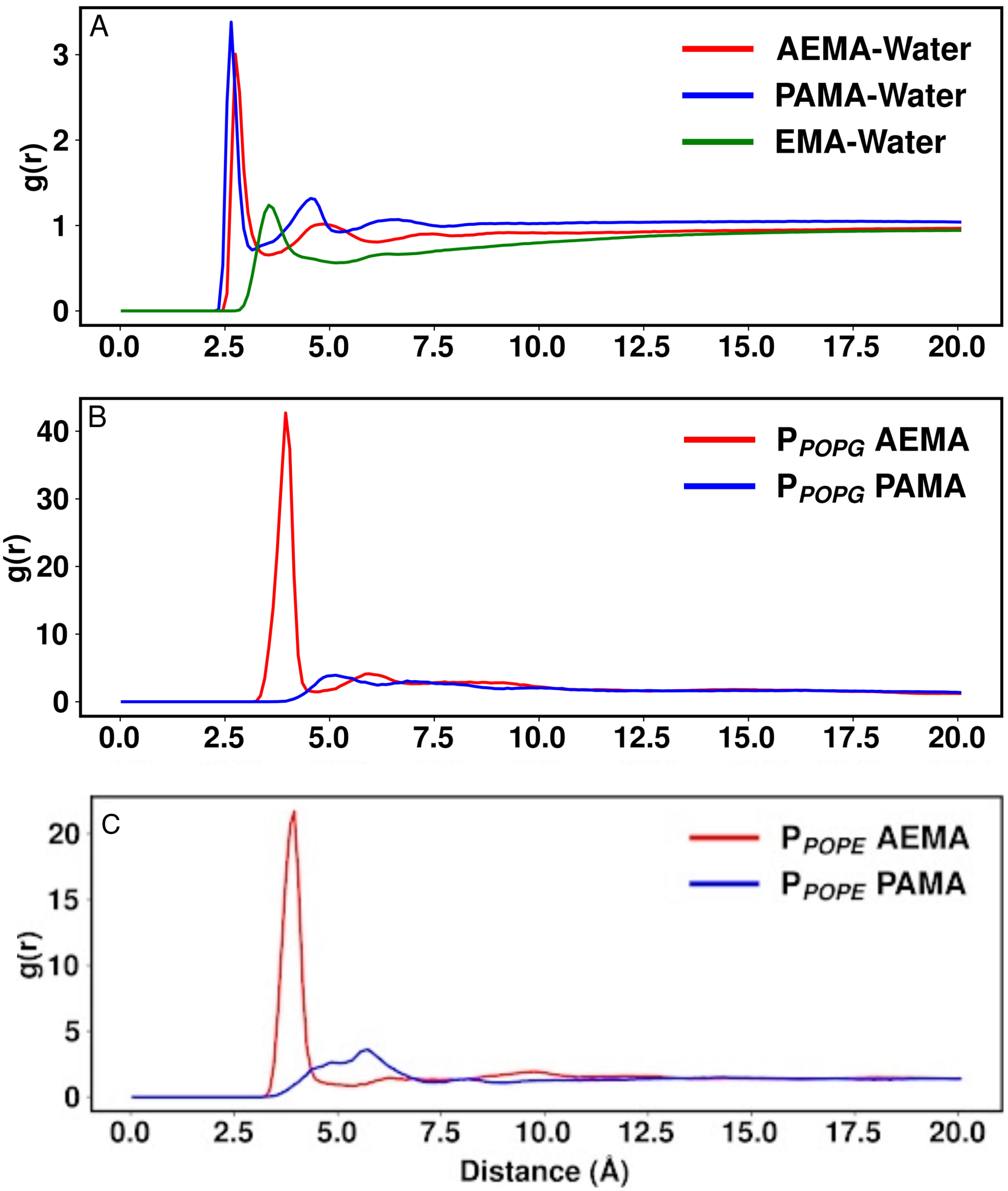


FIG. S6. Radial distribution function of AMPoly charged groups wrt POPE and POPG lipid charged groups.

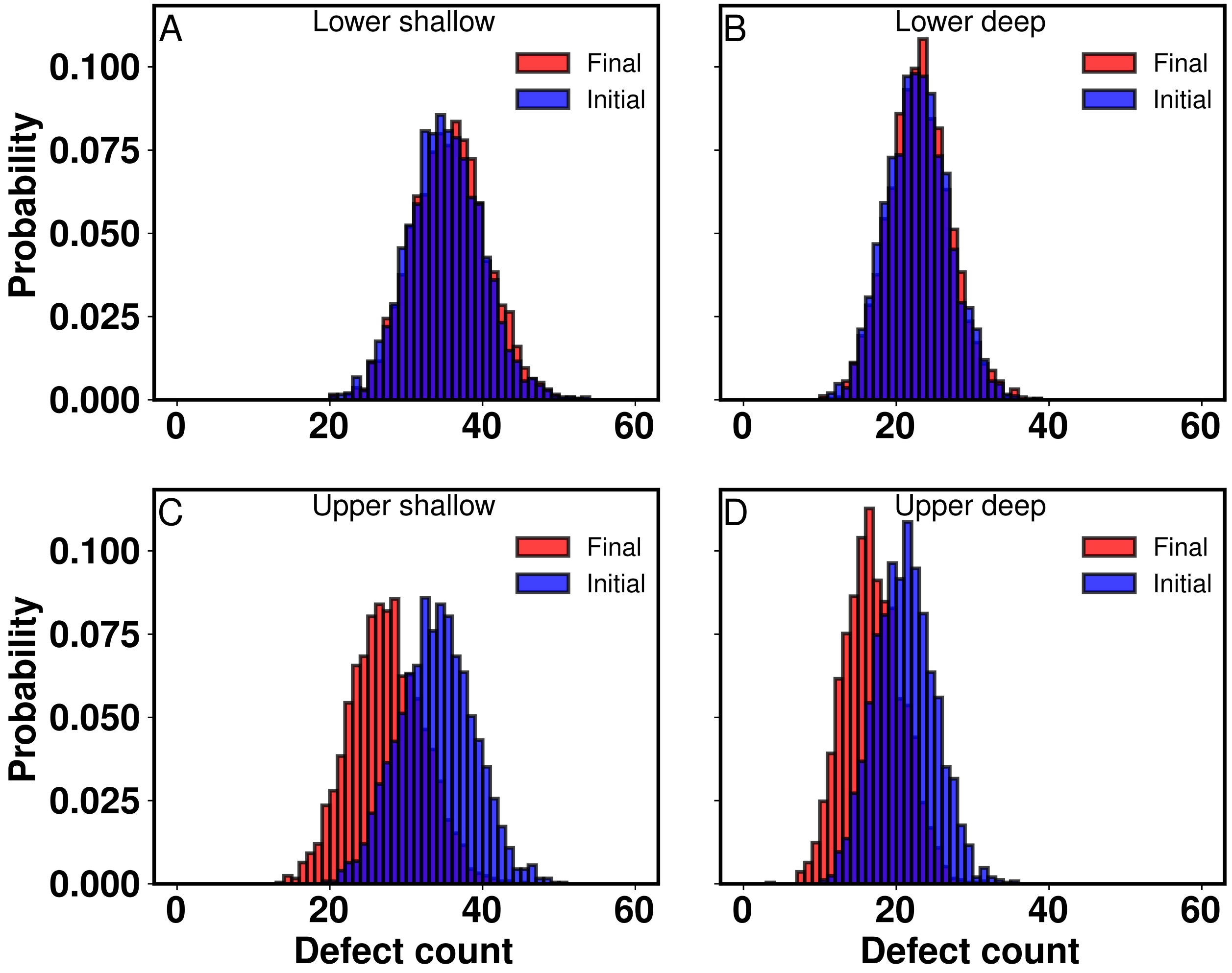


FIG. S7. Histogram of defect counts over the course of initial and final 300 ns of simulation. Presence of AMP in the upper leaflet reduces the number of both shallow and deep defects. No such change is seen in the lower leaflet.
